# A live-cell autophagy reporter reveals reversible vacuolation in naked mole-rat skin fibroblasts under lysosomal stress

**DOI:** 10.64898/2026.03.18.712644

**Authors:** Fangze Tong, Mateo P. Hoare, Laura J. Grundy, Filomena Gallo, Karin H. Müller, Ewan St. John Smith, Janet R. Kumita

## Abstract

Naked mole-rats (NMRs, *Heterocephalus glaber*) display unusual longevity and resistance to age-related decline, and accumulating evidence suggests that their autophagy–lysosome pathway (ALP) is regulated differently from that of conventional mammalian models. However, most studies in NMR cells have relied on static biochemical or ultrastructural readouts, leaving the dynamic organisation of autophagy in living cells poorly defined. Here, we establish a stable tandem fluorescent autophagy reporter in NMR skin fibroblasts using an mCherry–EGFP–LC3^NMR^ construct to enable live-cell, single-cell resolution analysis of ALP dynamics. Under basal conditions, NMR skin fibroblasts exhibit a greater abundance of LC3-positive structures than HeLa cells, together with a mixed population of autophagosomes and autolysosomes, indicating a distinct steady-state organisation of the ALP. Chloroquine (CQ)-induced lysosomal stress caused the expected accumulation of LC3-positive structures but also triggered the formation of large cytoplasmic vacuoles in NMR skin fibroblasts. Importantly, this vacuolation was not associated with acute cytotoxicity and progressively resolved following CQ removal, accompanied by reorganisation of LC3-positive compartments and recovery of lysosomal acidity. Electron microscopy showed that CQ-induced vacuoles are membrane-bound, containing internal material and co-existing with multiple ALP-related vesicular compartments. Primary NMR skin fibroblasts display a similar vacuolation phenotype, indicating that this response is not an artefact of immortalisation or reporter expression. Together, these findings establish a live-cell platform for analysing autophagy in NMR cells and identify a distinctive, reversible vacuolation response to lysosomal stress, consistent with dynamic remodelling of the lysosomal system within NMR skin fibroblasts.

## Introduction

Autophagy is a highly conserved catabolic process essential for maintaining intracellular homeostasis. It mediates the degradation and recycling of damaged proteins, dysfunctional organelles, and other cytoplasmic components through the lysosomal system [1]. Among the three major forms of autophagy (chaperone-mediated autophagy, microautophagy, and macroautophagy), macroautophagy (hereafter referred to as “autophagy”) is the predominant and most extensively studied pathway. It involves the sequestration of cytoplasmic material within double-membraned autophagosomes, that subsequently fuse with lysosomes to form autolysosomes, enabling cargo degradation and recycling [2]. Through this process, autophagy supports bioenergetic balance and preserves protein and organelle homeostasis, thereby playing a central role in cellular adaptation to stress and the maintenance of cellular health [3, 4]. Dysregulation of autophagy has been strongly implicated in ageing and in the pathogenesis of several diseases, including cancer, cardiovascular disorders, and neurodegenerative conditions [5–8].

Despite the well-established importance of autophagy in ageing and disease [5, 9, 10], therapeutic manipulation of this pathway has so far yielded limited clinical success [11, 12]. Therefore, one emerging paradigm is to investigate autophagy in naturally healthy ageing models, with the aim of understanding how long-term cellular homeostasis is maintained. The naked mole-rat (*Heterocephalus glaber*, NMR) represents a striking model for healthy ageing. NMRs possess a lifespan exceeding 35 years, approximately tenfold that of similarly sized mice [13, 14]. Beyond their exceptional longevity, NMRs display a natural resistance to age-associated pathologies, including cancer and neurodegenerative diseases [15, 16]. Although the molecular mechanisms underlying these unusual traits are not yet fully understood, accumulating evidence suggests that NMR cells undergo slower functional ageing and have altered proteostatic and autophagy-related homeostasis compared with conventional mammalian models [17–19]. The monitoring of autophagy in NMR cells has primarily been based on reporting end-point levels of autophagy-related reporters, such as LC3 I/II and beclin [18,20–22], using biochemical measurements such as western blotting and transcriptional analyses. In addition, transmission electron microscopy (TEM) analysis has described the appearance of cytosolic vacuoles in NMR skin fibroblasts following lysosomal inhibition [22]. Together, these findings suggest altered regulation of the autophagy–lysosome pathway (ALP) in NMRs; however, how these reported features manifest dynamically in living cells remains largely unexplored.

Autophagy is a highly dynamic and context-dependent process. Accordingly, static endpoint measurements, such as steady-state LC3-II or p62 levels, rarely provide an accurate reflection of autophagic flux, as they cannot distinguish between increased autophagosome biogenesis and impaired lysosomal clearance, two mechanistically distinct scenarios that may produce similar steady-state readouts [23–25]. To address these limitations, a range of live-cell assays for monitoring autophagic flux have been developed, including tandem fluorescent LC3 reporters (e.g., mCherry–EGFP–LC3), HiBiT-based systems, and other fluorescence-based approaches [2, 26–28]. These tools have been widely applied in mammalian cell models to resolve autophagosome formation, maturation, and lysosomal degradation at single-cell resolution, thereby substantially advancing our understanding of autophagy regulation in ageing and disease contexts. However, despite their broad utility, such live-cell, flux-resolved assays have not been systematically applied to NMR cells, leaving a critical gap in knowledge regarding the dynamic regulation of autophagic flux and its response to perturbations within the ALP.

To overcome this, we have established a stable autophagy reporter cell line in NMR skin fibroblasts using a tandem fluorescent mCherry–EGFP–LC3^NMR^ construct, enabling real-time visualisation of autophagic flux at single-cell resolution by differentially reporting on autophagosome and autolysosome/lysosome populations. Using this platform, we systematically perturbed distinct regulatory steps of the ALP, including mTOR signalling, lysosomal acidification, and late-stage autophagosome maturation, and integrated live-cell imaging with ultrastructure analyses to interrogate flux regulation under the different stresses [29–31]. This experimental framework allows us to characterise autophagy dynamics, vacuolar remodelling, and recovery following lysosomal stress in NMR skin fibroblasts. Using this platform, we were able to resolve chloroquine-induced vacuolation in live NMR cells and track the time-dependent morphological changes that occur during lysosomal inhibition and subsequent recovery, enabling us to assess whether this vacuolation is reversible and to capture dynamic transitions not accessible with static endpoint measurements. To validate these fluorescence-based observations, we further employed electron microscopy techniques to confirm the ultrastructural characteristics of the ALP-associated compartments. Together, this integrated framework provides a dynamic reinterpretation of previously reported autophagy phenotypes in NMR cells.

## Results

### Validation of a tandem fluorescent autophagy reporter in NMR skin fibroblasts

Site-directed mutagenesis was used to change three amino acids in the mammalian cell reporter plasmid expressing mCherry–EGFP–LC3 (Addgene #123230) to create the NMR version of LC3 (E105G, M121R, K122G) (mCherry–EGFP–LC3^NMR^). Following transfection and antibiotic selection, multiple single-cell–derived colonies were isolated to establish stable NMR skin fibroblast lines expressing the tandem fluorescent mCherry–EGFP–LC3^NMR^ reporter. These NMR cells are skin fibroblasts immortalised by SV40 transfection that have been previously characterised [32]. To verify the expression and integrity of the fusion protein, selected colonies were analysed by western blotting (WB) using antibodies against LC3 and EGFP (**Figure 1A, B**). Both antibodies detected two bands (non-lipidated and lipidated LC3 isoforms) at approximately 80–82 kDa, consistent with the predicted molecular mass of the full-length mCherry–EGFP–LC3^NMR^ fusion protein.

**Figure 1:**
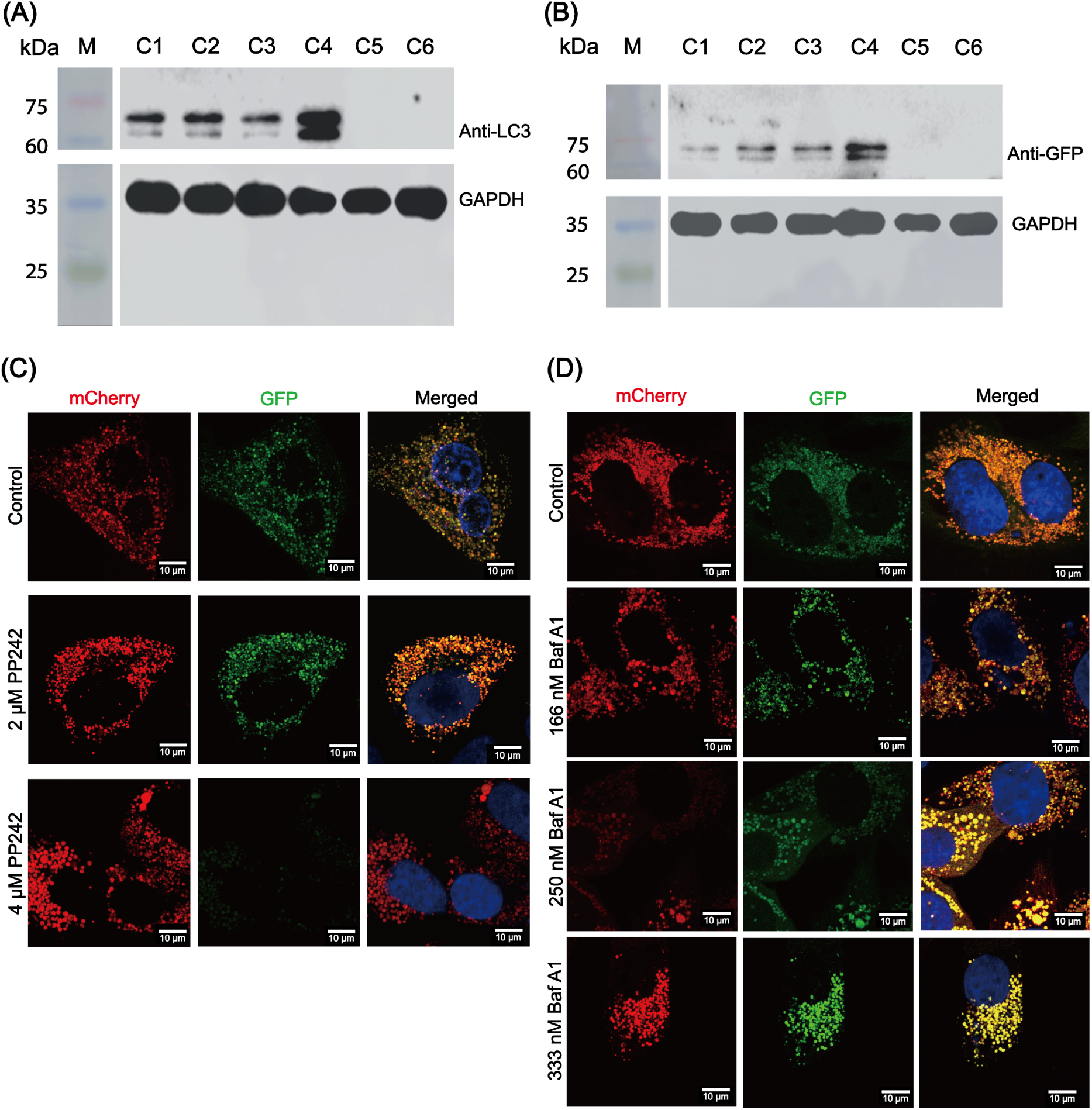
Characterisation of the mCherry–EGFP–LC3^NMR^ reporter NMR skin fibroblast cell line. Western blot validation of mCherry–EGFP–LC3^NMR^ stable over-expression in NMR skin fibroblast lines. Cell lysates from single-cell–derived colonies were probed with antibodies against LC3 (**A**) and EGFP (**B**). Both antibodies detected doublet bands at approximately 80–82 kDa, consistent with the predicted molecular mass of the mCherry–EGFP–LC3^NMR^ fusion protein in the non-lipidated (LC3-I; top band) and lipidated (LC3-II, bottom band) forms. Markers are shown in lane M and individual colonies are labelled C1-C6. **C**) mTOR inhibition by PP242 enhances autophagic flux in NMR skin fibroblasts. Representative live-cell confocal images of mCherry–EGFP–LC3^NMR^ expressing cells, either untreated or treated with PP242 (2 μM or 4 μM) for 24 h. PP242 treatment induces a shift from the appearance of autophagosomes (mCherry⁺/EGFP⁺ (yellow)) to autolysosomes (mCherry⁺/EGFP⁻ (red)). Scale bar is 10 μm. **D)** Bafilomycin A1 (Baf A1) impairs autophagic flux in NMR skin fibroblasts. Representative live-cell confocal images of mCherry–EGFP–LC3^NMR^ expressing cells, either untreated or treated with Baf A1 (166 nM, 250 nM, or 333 nM) for 24 h. Baf A1 treatment results in accumulation of mCherry⁺/EGFP⁺ (yellow) puncta, consistent with impaired autophagic flux. In **C)** and **D)** the nucleus is stained with Hoescht (blue).

Among the colonies positive for both LC3 and EGFP signals, a clone exhibiting intermediate expression levels (colony 2) was selected for all subsequent experiments to minimise potential artefacts associated with overexpression. To functionally validate the reporter, we next examined its response to pharmacological modulators of autophagy using fluorescent microscopy. Treatment with PP242, an mTOR inhibitor, induced a dose-dependent shift from mixed-colour puncta towards predominantly mCherry⁺/EGFP^-^ (red) LC3 puncta, consistent with increased autophagic flux and efficient progression of LC3-labelled structures to acidic compartments (**Figure 1C**) [26, 29, 33]. In contrast, dose-dependent inhibition of lysosomal acidification with bafilomycin A1 (Baf A1) led to a pronounced accumulation of mCherry⁺/EGFP⁺ LC3 (yellow) puncta, consistent with impaired autophagic flux and accumulation of autophagosomes upstream of lysosomal degradation (**Figure 1D**) [26, 30].

Together, these data confirm stable expression of the mCherry–EGFP–LC3^NMR^ fusion protein in NMR skin fibroblasts and demonstrate that this tandem fluorescent reporter faithfully monitors changes in autophagic flux in response to both autophagy induction and lysosomal inhibition.

### Basal level autophagic profiles and lysosomal organisation differ between NMR skin fibroblasts and HeLa cells

We compared untreated, basal level culture conditions for tandem fluorescent mCherry–EGFP–LC3 reporter lines in NMR skin fibroblasts and human HeLa cells (**Figure 2**). In the HeLa reporter cells, very few puncta are observed, but those LC3-positive puncta present were mCherry⁺/EGFP⁻ (red), corresponding to acidified autolysosomal structures (**Figure 2A**). In contrast, the NMR skin fibroblast reporter cells display significantly more puncta, which is consistent with previous reports suggesting altered basal autophagy-related activity in NMR cells [18], and they are a mixed population of mCherry⁺/EGFP⁻ (red) and mCherry⁺/EGFP⁺ (yellow) LC3 puncta, suggesting differences in basal autophagic organisation within the two cell types.

**Figure 2:**
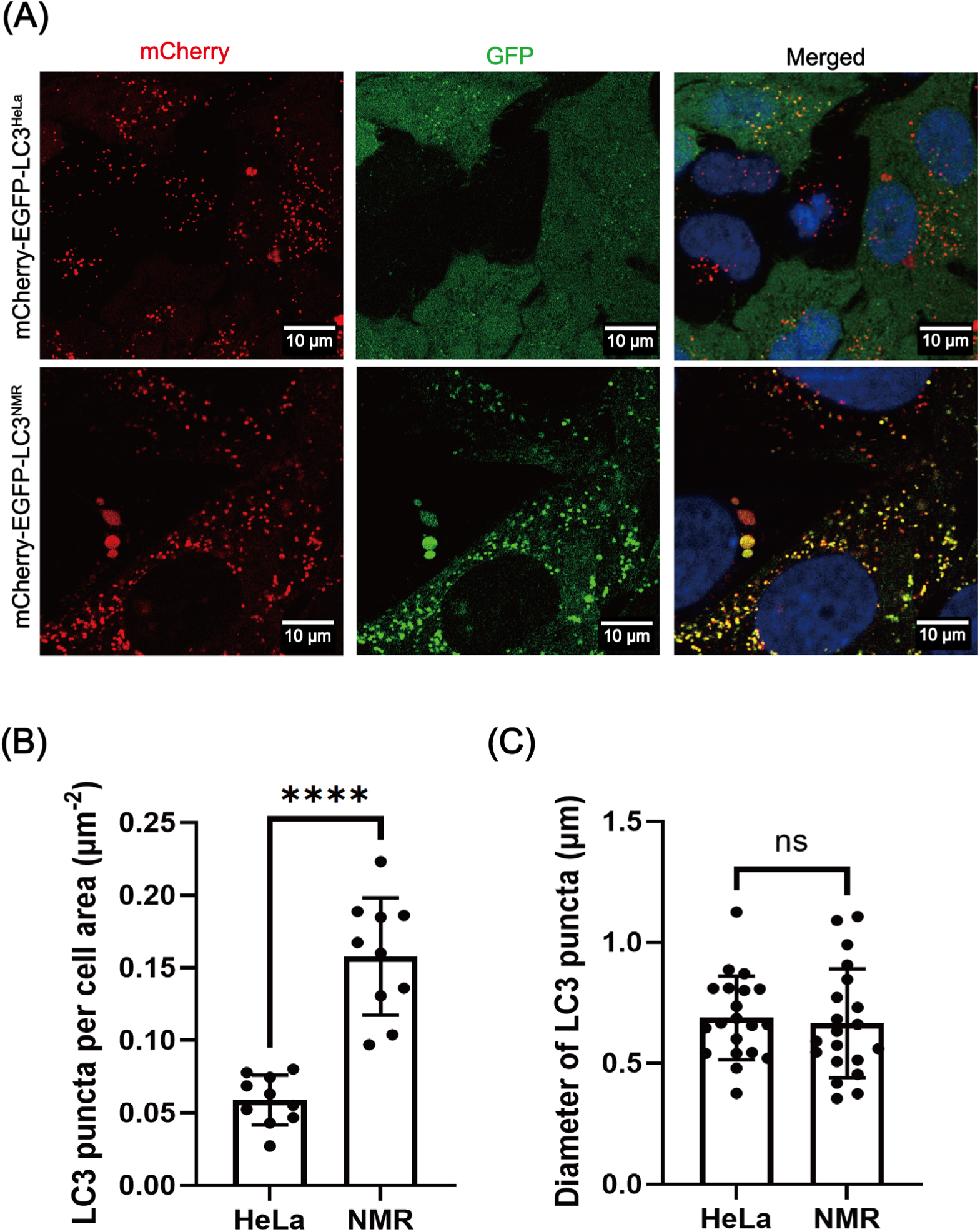
Tandem fluorescent autophagy reporter reveals distinct basal autophagic profiles in HeLa cells and NMR skin fibroblasts. **A**) Representative live-cell confocal images of HeLa cells and NMR skin fibroblasts stably expressing mCherry–EGFP–LC3 under basal-level culture conditions. The nucleus is stained with Hoescht (blue). Few puncta are present in the HeLa cells, and those that are correspond to mCherry^+^/EGFP^-^ (red). In contrast, NMR skin fibroblasts display more puncta with a mixture of mCherry^+^/EGFP^-^ (red) and mCherry^+^/EGFP^+^ (yellow). **B**) Quantification of LC3 puncta density in HeLa and NMR skin fibroblasts, normalised to cell area (puncta per μm²). Ten individual cells per cell line were analysed, sampled from four independent fields of view. Statistical significance was assessed using a two-tailed unpaired t-test. ****P < 0.0001. **C**) Quantification of LC3 puncta diameter in HeLa cells and NMR skin fibroblasts. Twenty individual LC3 puncta per cell line were analysed, sampled from four independent fields of view. Statistical analysis was performed using a two-tailed unpaired t-test. n.s. - not significant.

To further characterise these differences, we quantified LC3 puncta density normalised to cell area, as well as puncta size, irrespective of whether they were autophagosomes or autolysosomes. NMR skin fibroblasts exhibited a significantly higher density of LC3-positive puncta compared with HeLa cells (p < 0.0001; **Figure 2B**). In contrast, the average diameter of LC3 puncta did not differ significantly between NMR skin fibroblasts and HeLa cells (**Figure 2C**). Together, these data indicate that, relative to the HeLa cells, NMR skin fibroblasts display an increased abundance of LC3-labelled autophagic structures in basal-level conditions, without an accompanying change in puncta size.

### Chloroquine-induced lysosomal stress elicits pronounced vacuolation in NMR skin fibroblasts

A persistent challenge in autophagy research is distinguishing enhanced autophagosome biogenesis from impaired degradation [19]. To explicitly test whether NMR skin fibroblasts regulate autophagic flux differently under conditions where degradation becomes limiting, we used chloroquine (CQ), a lysosomotropic agent that impairs lysosomal acidification and autophagic degradation, to impose tuneable lysosomal stress [17, 27]. This manipulation imposed a defined constraint on the ALP, enabling assessment of how NMR skin fibroblasts accumulate and process LC3-positive structures when lysosomal clearance is compromised. When NMR reporter skin fibroblasts were treated with different concentrations of CQ for 4 h, a progressive accumulation of mCherry⁺/EGFP⁺ (yellow) LC3 puncta was observed (**Figure 3**), consistent with impaired lysosomal degradation. Interestingly, treatment with increasing CQ also induces the formation of large cytoplasmic vacuoles, which were readily apparent in live-cell imaging (**Figure 3A**). To further characterise the autophagic response under these conditions, we measured LC3 levels by western blotting. CQ treatment resulted in the expected accumulation of LC3-II (**Figure 3B**), with significantly increased LC3-II/LC3-I ratios compared with untreated controls (**Figure 3C**). Together, these data show that NMR skin fibroblasts respond to lysosomal stress with LC3-II accumulation and increased abundance of LC3-positive structures, consistent with impaired autophagic degradation. Notably, lysosomal stress induced by CQ triggers a pronounced vacuolation phenotype in NMR skin fibroblasts, suggesting distinctive cellular remodelling responses under conditions of compromised lysosomal clearance.

**Figure 3:**
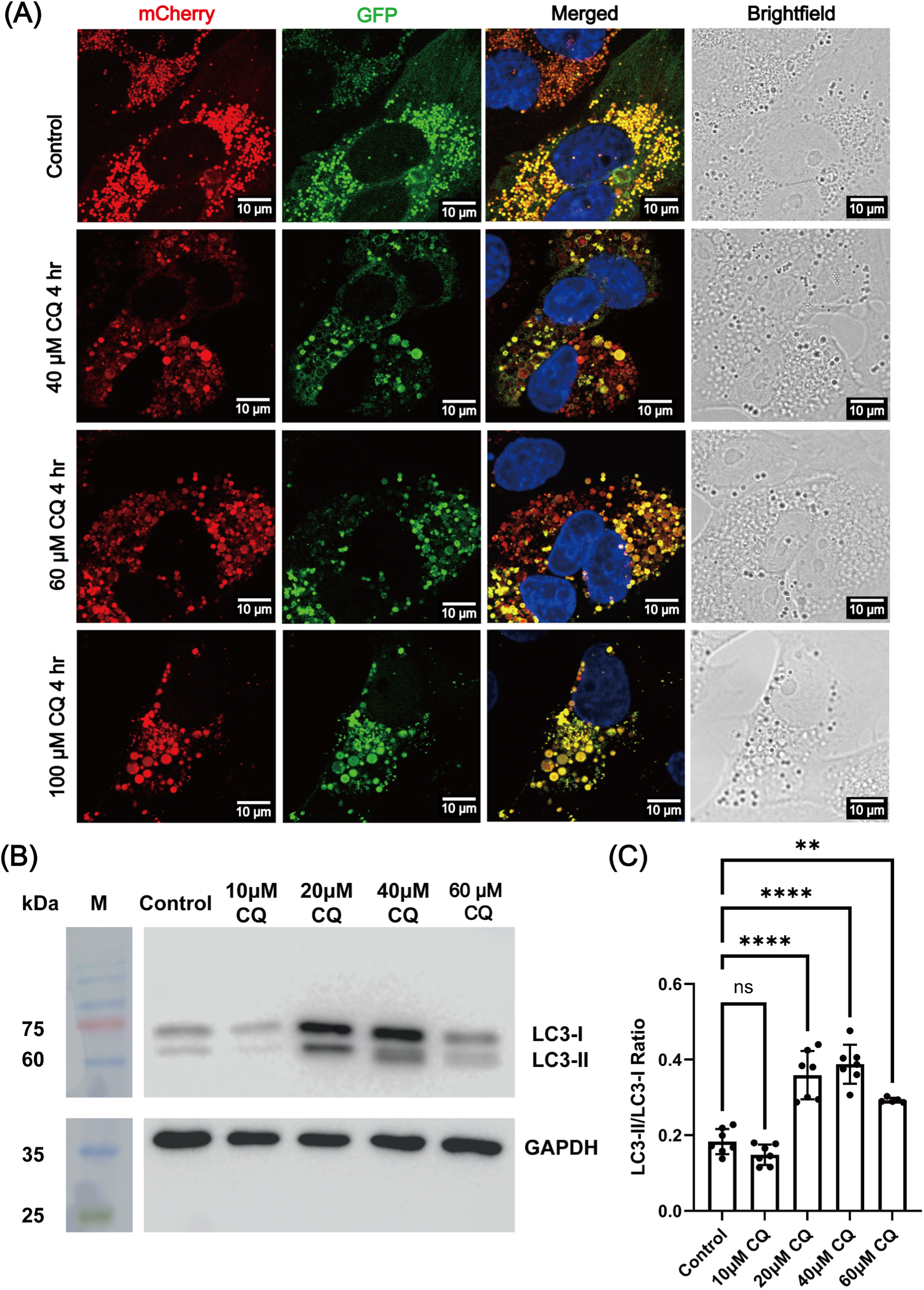
Chloroquine treatment inhibits autophagic flux and induces vacuolation in the mCherry–EGFP–LC3^NMR^ reporter NMR skin fibroblast cell line. **A)** Representative live-cell confocal images of mCherry–EGFP–LC3^NMR^ expression in cells either untreated or treated with increasing concentrations of CQ (40, 60 and 100 μM) for 4 h. The nucleus is stained with Hoescht (blue). CQ-treated cells exhibited predominantly mCherry⁺/EGFP⁺ (yellow) puncta, consistent with impaired lysosomal degradation. Brightfield panels show the appearance of the cytoplasmic vacuolation with increasing CQ concentrations. **B**) WB analysis of mCherry–EGFP–LC3^NMR^ protein levels from cells either untreated or treated with increasing concentrations of CQ (10 μM, 20 μM, 40 μM, or 60 μM) for 24 h. CQ treatment resulted in an accumulation of LC3-II relative to LC3-I. (**C**) Quantification of LC3-II/LC3-I ratios derived from densitometric analysis. CQ treatment significantly increases LC3-II accumulation relative to controls. **P < 0.01, ***P < 0.001, ****P < 0.0001.

### Vacuoles induced by CQ in NMR skin fibroblasts are decorated with autophagy-related markers

We next sought to further characterise the nature of the large cytoplasmic vacuoles induced by CQ treatment. Because CQ primarily perturbs lysosomal function, we asked whether the observed vacuoles were associated with the ALP (i.e. exhibiting autophagic or lysosomal features), or whether they instead represent non-specific cytoplasmic swelling. To systematically assess the molecular composition and organisation of CQ-induced vacuoles in NMR skin fibroblasts, we performed time-dependent live-cell imaging, using the mCherry–EGFP–LC3^NMR^ expressing NMR cell-line treated with 40 μM CQ for 16 or 24 h. This analysis revealed features that were distinct from those observed under other autophagy-modulating conditions. After 16 h of CQ treatment, both mCherry⁺/EGFP⁻ (red) and mCherry⁺/EGFP⁺ (yellow) LC3 puncta were present (**Figure 4**). In addition, prominent LC3-positive vacuolar structures became evident, including EGFP⁺/mCherry⁺ (yellow) vacuoles enclosing distinct red puncta (white arrow, **Figure 4**) and discrete ring-like EGFP⁺/mCherry⁺ structures (pink arrow, **Figure 4**). These LC3-labelled ring structures appear to be associated with the surface of cytoplasmic vacuoles. Following 24 h of CQ treatment, the abundance of LC3-labelled vacuoles increased further (**Figure 4**, lower panel). These findings show that the CQ-induced vacuoles in NMR skin fibroblasts are closely associated with the ALP and are decorated by autophagy-related markers, rather than representing non-specific cytoplasmic swelling. Interestingly, LC3 labelling appeared progressively and did not always coincident with the earliest stages of vacuole formation. This temporal separation suggests that LC3 recruitment may occur after the initial vacuolar remodelling triggered by lysosomal stress.

**Figure 4:**
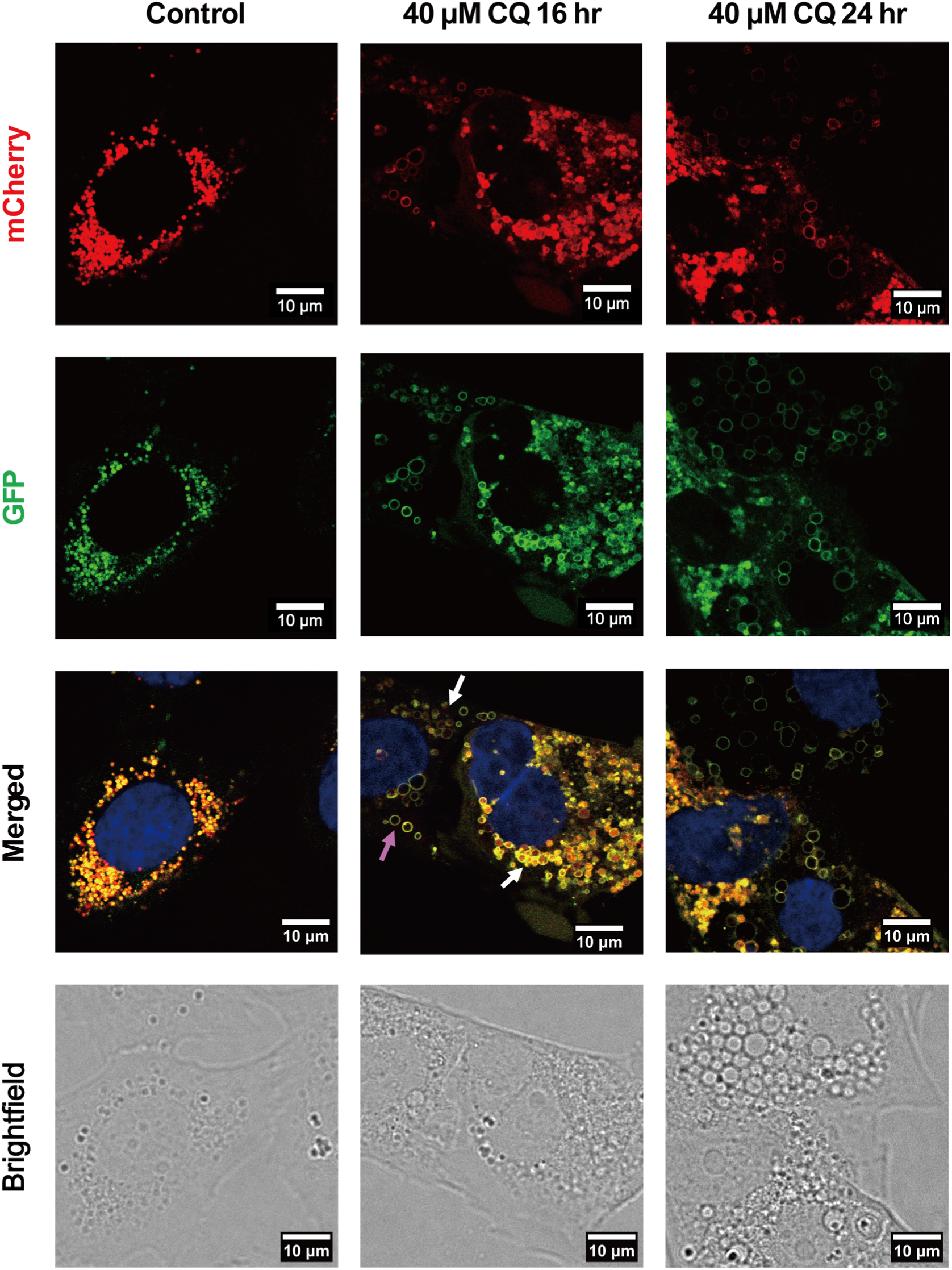
CQ-induced vacuoles are associated with LC3-positive structures in the mCherry–EGFP–LC3^NMR^ reporter NMR skin fibroblast cell line. Representative live-cell confocal images monitoring mCherry–EGFP–LC3^NMR^ following CQ treatment. The nucleus is stained with Hoescht (blue). Cells were treated with 40 μM CQ for 16 h or 24 h, as indicated. In the control (basal level), LC3-positive puncta are observed as mCherry⁺/EGFP⁻ (red) or mCherry⁺/EGFP⁺ (yellow) structures. After 16 h of CQ treatment, additional LC3-positive structures emerge, including mCherry⁺/EGFP⁺ ring-like structures associated with the surface of large cytoplasmic vacuoles (pink arrow) and mCherry⁺/EGFP⁺ ring-like structures enclosing red puncta (white arrow). Following 24 h of CQ treatment, the abundance of LC3-labelled vacuoles increases further.

### CQ-induced vacuoles in NMR skin fibroblasts are not associated with acute cell death and are reversible

Given the pronounced and dynamic nature of CQ-induced vacuolation, we next examined the relationship between this phenotype and cell viability. To determine whether CQ treatment and the associated vacuolation were accompanied by cytotoxicity, a lactate dehydrogenase (LDH)-based cytotoxicity assay was performed on NMR skin fibroblasts (**Supplementary Figure 1**). Even at the highest CQ concentration tested (100 µM), where vacuolation is most pronounced, NMR skin fibroblasts show minimal cytotoxicity. Notably, under the same conditions, mouse skin fibroblasts exhibit significantly higher cytotoxicity, supporting that CQ-induced vacuolation in NMR skin fibroblasts is not simply a secondary consequence of acute cell death [31, 34, 35].

Using time-dependent live imaging experiments, we next looked at whether the vacuoles were reversible after the removal of lysosomal stress by treating the NMR reporter skin fibroblasts with 40 μM CQ for 24 h, followed by removing CQ and allowing the cells to recover in drug-free media. Images were taken at 4 h and 24 h post-CQ removal to monitor vacuole resolution and autophagic organisation (**Figure 5**). After 4 h of recovery from CQ treatment, some vacuoles remained but most large vacuoles disappeared and LC3-positive ring-like structures, previously associated with vacuolar structures, became fewer. At 24 h post-CQ removal, LC3-labelled vacuoles were no longer present and the majority of LC3 puncta exhibited a distribution comparable to untreated control skin fibroblasts, consistent with resumed autophagic flux, as indicated by the appearance of small, mCherry^+^/EGFP^-^ (red) puncta indicative of autolysosome and lysosome formation (**Figure 5**, upper and lower panels). As a control for recovery after CQ treatment, these experiments were also performed on mCherry–EGFP–LC3^WT^ expressing HeLa cells (**Supplementary Figure 2**), where CQ treatment results in the expected accumulation of mCherry⁺/EGFP⁺ (yellow) LC3 puncta (not vacuoles); however, in contrast to the NMR skin fibroblasts, removal of CQ did not result in efficient recovery under identical exposure and recovery conditions, with the majority of LC3 puncta in HeLa cells remained mCherry⁺/EGFP⁺ (yellow) after 24 h of recovery.

**Figure 5.**
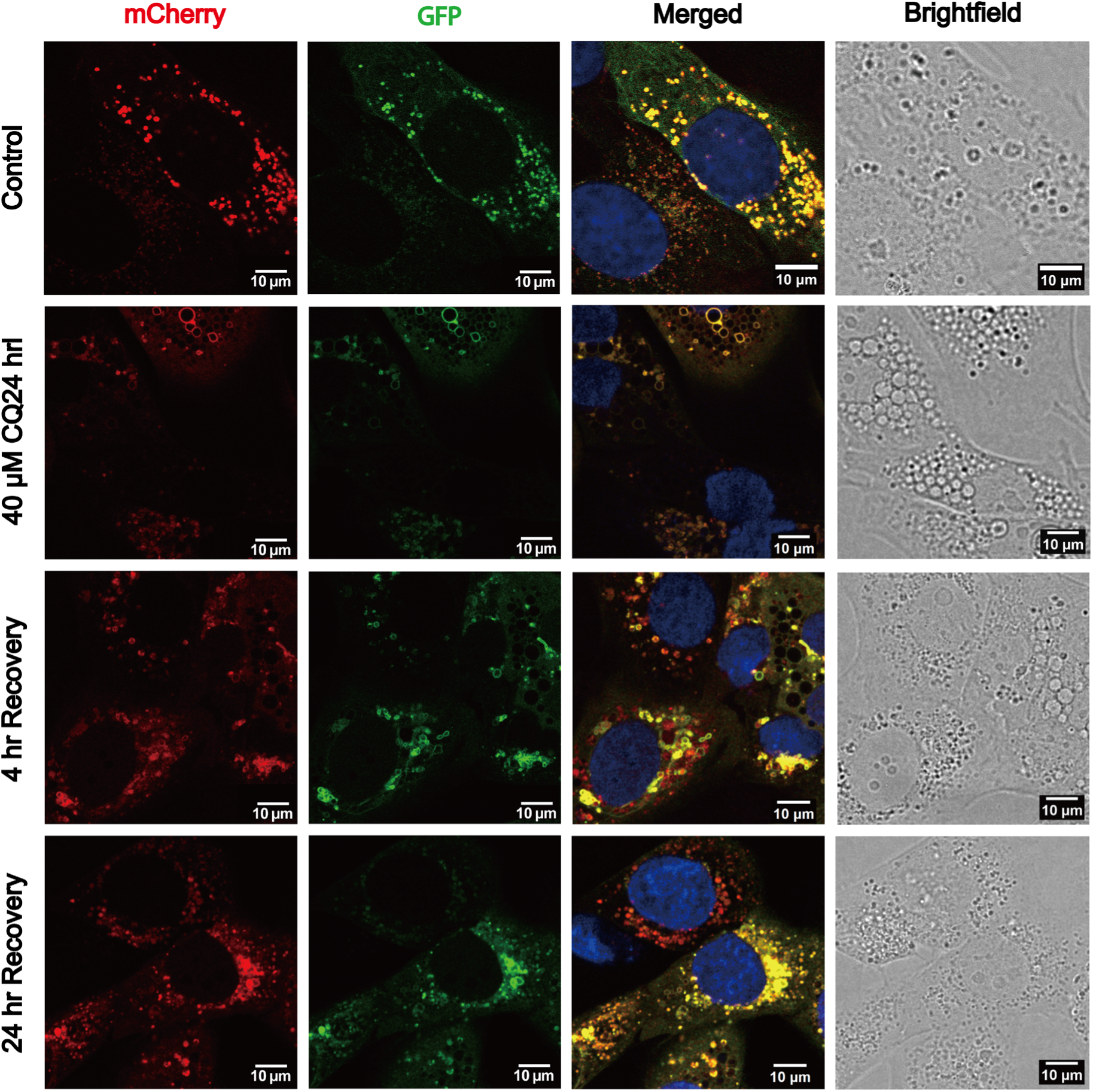
Recovery of autophagic organisation in the mCherry–EGFP–LC3^NMR^ reporter NMR skin fibroblast cell line following CQ removal. Representative live-cell confocal images of mCherry–EGFP–LC3^NMR^ expression in cells left untreated, treated with CQ (40 μM) for 24 h, or allowed to recover for 4 h or 24 h following CQ removal, as indicated. The nucleus is stained with Hoescht (blue). After 24 h of CQ treatment, LC3-positive structures predominantly appear as mCherry^+^/EGFP^+^ (yellow) puncta and LC3-decorated ring-like vacuolar structures. Following the removal of CQ from the media, progressive reorganisation of LC3-labelled structures is observed. At 4 h of recovery, smaller mCherry^+^/EGFP^-^ LC3 puncta and mCherry^+^/EGFP^+^ structures are frequently observed, and ring-like structures are less apparent. By 24 h of recovery, LC3 labelling is no longer associated with vacuoles, and the majority of LC3 puncta exhibit a distribution comparable to untreated control cells.

From the analysis of the LC3-reporter, it appears that removing CQ leads to a return to basal-level autophagy, suggested by the presence of both autophagosomes (mCherry^+^/EGFP^+^) and autolysosome-like structures (mCherry^+^/EGFP^-^) (**Figure 5**). However, to understand if the recovered morphology recovered functional autolysosome attributes, we assessed the acidity of CQ-induced vacuoles in NMR skin fibroblasts (not overexpressing mCherry–EGFP–LC3^NMR^) using LysoTracker Red, a fluorescent probe that selectively accumulates in acidic compartments [36]. As validation controls, NMR skin fibroblasts were treated with Baf A1 (300 nM) or PP242 (4 μM). As expected, Baf A1 abolishes LysoTracker Red staining, whereas PP242 treatment enhances LysoTracker Red signal intensity (**Supplementary Figure 3**). Lysotracker Red signals were monitored in NMR skin fibroblasts that were untreated, treated with CQ (100 μM) for 24 h, or allowed to recover for 4 h or 24 h following CQ removal (**Figure 6**). LysoTracker Red signal appeared overall more prominent in CQ-treated cells; however, its distribution was heterogeneous, with only a subset of vacuoles showing clear LysoTracker association, while many vacuoles lacked detectable LysoTracker signal (**Figure 6**). During recovery, the vacuolation subsided, and an increase in puncta co-localising with LysoTracker Red signal was observed (green arrows), consistent with progressive reacquisition of acidic properties in these vacuolar compartments.

**Figure 6.**
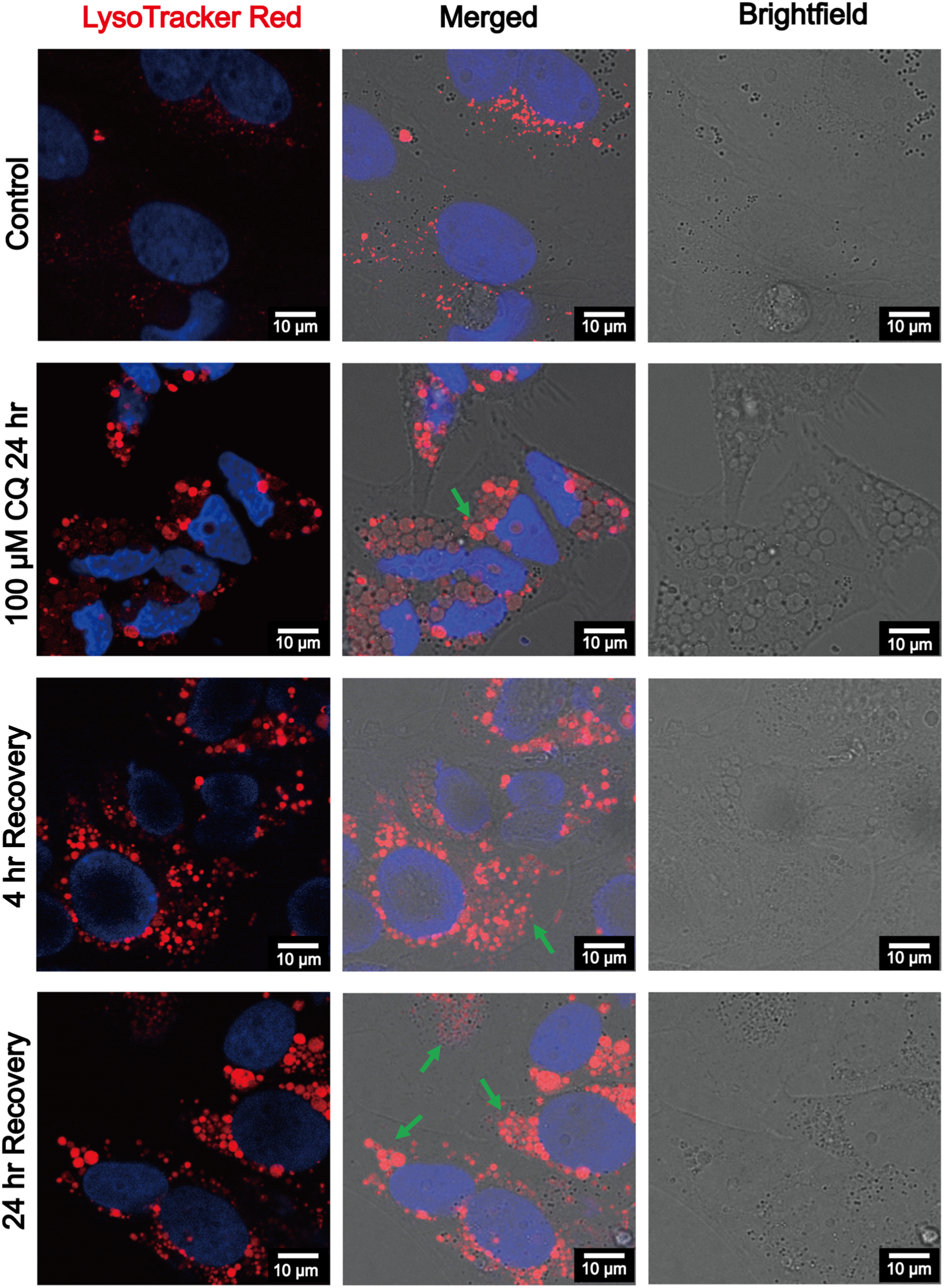
LysoTracker Red staining during recovery from CQ treatment in non-transfected NMR skin fibroblasts. Representative live-cell confocal images of immortalised NMR skin fibroblasts stained with LysoTracker Red and Hoescht (blue – nucleus). Cells were either untreated, treated with CQ (100 μM) for 24 h, or allowed to recover for 4 h or 24 h following CQ removal, as indicated. In CQ-treated cells, LysoTracker Red staining is not uniformly associated with all vacuoles. During recovery, LysoTracker-positive puncta and vacuolar structures become more apparent (green arrows), consistent with progressive reacquisition of acidic properties.

### Electron microscopy reveals the ultrastructural features of CQ-induced vacuoles in NMR skin fibroblasts

Having established that CQ-induced vacuoles are associated with ALP, we next sought to define their ultrastructural characteristics to corroborate our fluorescence microscopy imaging. Because the presence of both mCherry⁺/EGFP⁺ and mCherry⁺/EGFP^-^ LC3 signals complicates the interpretation based on fluorescence imaging alone, we employed electron microscopy techniques to further characterise the morphology and organisation of these vacuoles. Scanning electron microscopy (SEM) revealed that CQ-induced vacuoles in NMR skin fibroblasts appear non-empty, in contrast to the apparently empty luminal spaces observed by live-cell fluorescence imaging (**Figure 7A**, middle panel). Following 24 h of recovery after CQ removal, most of the vacuolation has disappeared (**Figure 7A**, bottom panel), consistent with the live-imaging observations (**Figure 5**). Notably, although vacuolar structures appear to reorganise into LC3 puncta resembling those observed in untreated cells, SEM analysis indicated that these structures became more electron-dense during recovery (**Figure 7A**, lower panel).

**Figure 7.**
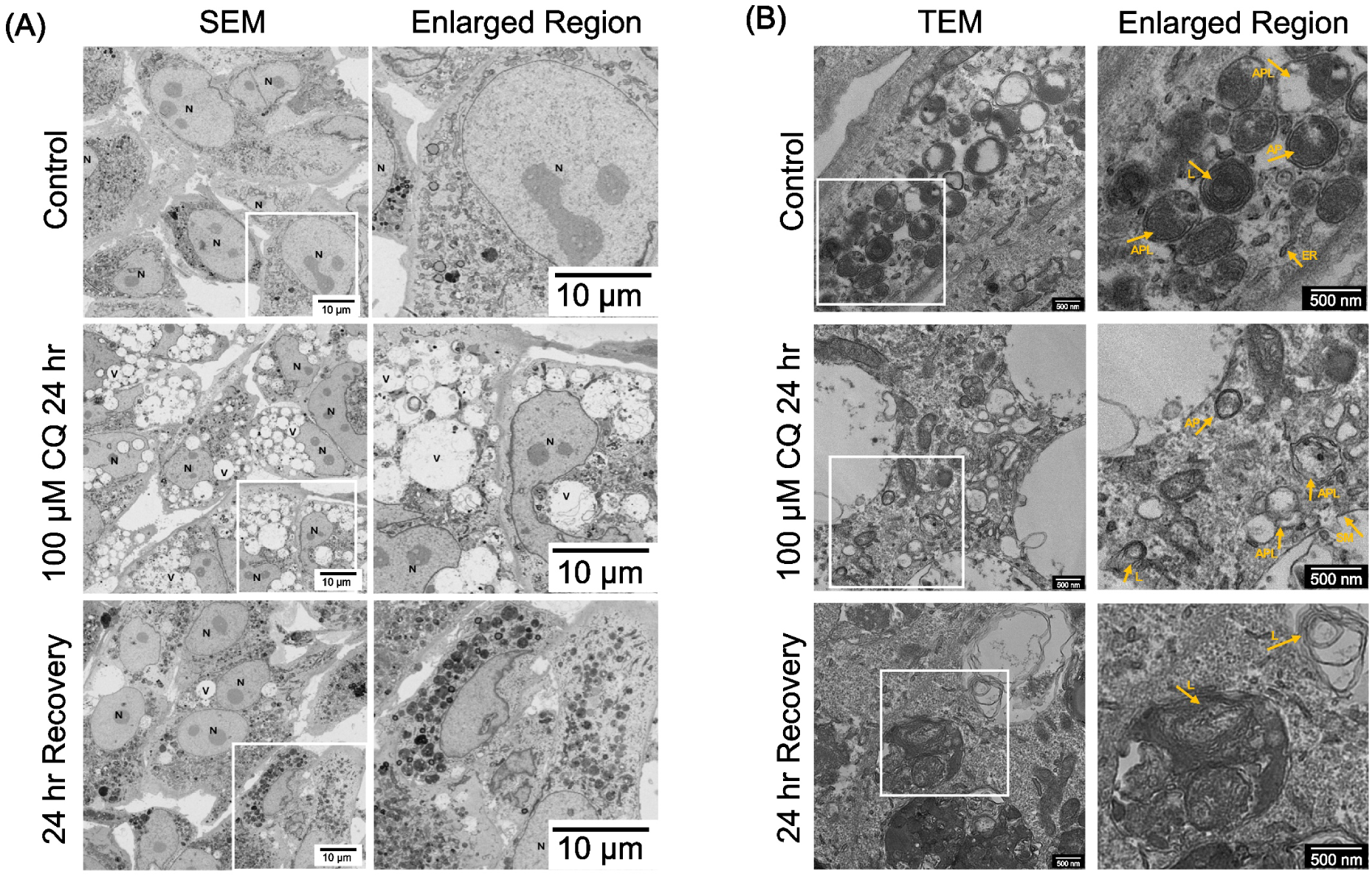
Electron microscopy imaging of CQ-induced vacuolation and recovery in NMR skin fibroblasts. **A)** Representative Scanning Electron Microscopy (SEM) images of immortalised NMR skin fibroblasts that were either untreated, treated with CQ (100 μM) for 24 h, or allowed to recover for 24 h following CQ removal. SEM analysis shows that CQ-induced vacuolar structures are not empty. Following 24 h of recovery after CQ removal, NMR skin fibroblasts exhibit an increased abundance of electron-dense structures, consistent with ultrastructural reorganisation during recovery. N – nucleus, V – vacuoles. **B)** Representative transmission electron microscopy (TEM) images of NMR skin fibroblasts under the same conditions as **A)**. A range of ALP–related vesicular structures are identified, including autophagosomes (AP), lysosomes (L), autophagosome-lysosome fusion intermediates (APL) and endoplasmic reticulum (ER). In addition, following treatment with 100 μM CQ for 24 h (middle panel), large vacuolar structures appear to be enclosed by single membranes (SM) and multiple autophagy-related vesicular structures are present in the cytoplasm between vacuoles.

Using transmission electron microscopy (TEM), the details of different structures involved in the ALP pathway can be resolved [23, 37]. In untreated NMR skin fibroblasts, multiple ALP-related vesicular structures are identified (**Figure 7B**, upper panel). Following CQ treatment, vacuoles appear as large vesicular structures enclosed by a single membrane (**Figure 7B**, middle panel). In addition, multiple autophagy-related vesicles are observed in the cytoplasm between vacuoles, including structures consistent with autophagosomes and autolysosomes (**Figure 7B**, middle panel). Following recovery from CQ treatment, TEM images indicate that vacuoles reorganise into smaller, lysosome-like structures characterised by increased electron density and membranous organisation (**Figure 7B**, lower panel), consistent with our live-cell imaging and LysoTracker Red staining (**Figure 6**).

### CQ-induced vacuolation was also evident in primary NMR skin fibroblasts

The mCherry–EGFP–LC3^NMR^ cell line was created with immortalised NMR skin fibroblasts, and therefore, to ensure the CQ-induced vacuolation is not an artefact, we tested CQ treatment on primary NMR skin fibroblasts. Primary cells were treated with CQ (20 μM or 100 μM) for 24 h and analysed by immunofluorescence confocal imaging (probing for LC3). Consistent with the observations in our mCherry–EGFP–LC3^NMR^ cell line and regular immortalised NMR skin fibroblasts, CQ treatment of primary NMR skin fibroblasts results in the formation of prominent cytoplasmic vacuoles (**Figure 8**). These vacuoles are readily detectable by brightfield imaging and colocalised with LC3, indicating engagement of the ALP. Notably, although the vacuolation phenotype in primary skin fibroblasts was qualitatively similar to that observed in immortalised skin fibroblasts, vacuole architecture and abundance in primary NMR skin fibroblasts appear to exhibit greater heterogeneity. This divergence is consistent with known phenotypic differences between primary cells and immortalised cell lines [38, 39]. Together, these data demonstrate that CQ-induced vacuolation is not an artefact of immortalisation and transfection and is retained in primary skin fibroblasts.

**Figure 8.**
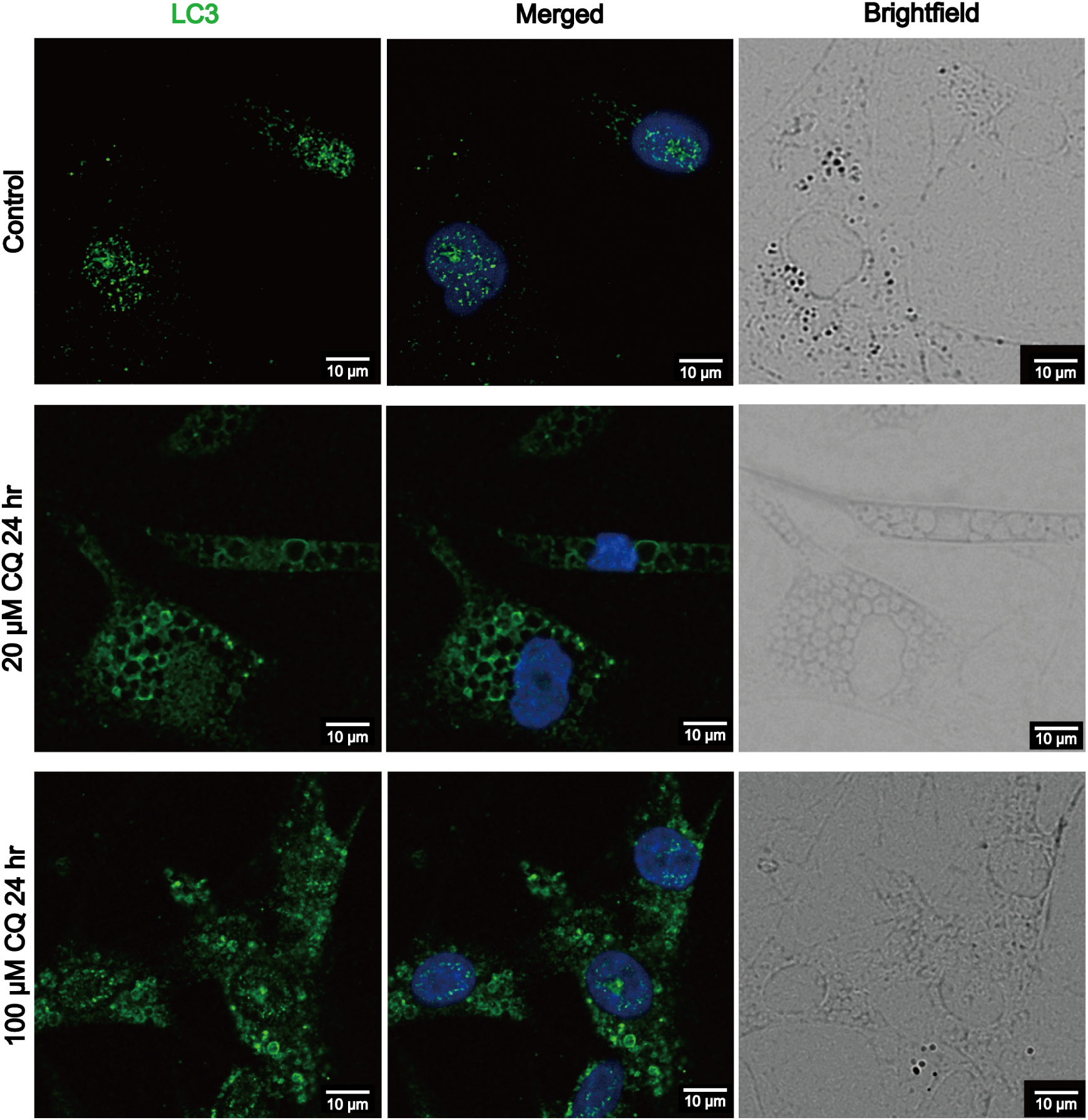
CQ-induced vacuolation in primary NMR skin fibroblasts. Representative immunofluorescence images of primary NMR skin fibroblasts that were either untreated or treated with CQ (20 μM and 100 μM) for 24 h. CQ treatment induces the formation of prominent cytoplasmic vacuoles that are readily visible by brightfield imaging. Vacuolar structures are frequently associated with LC3-positive signals (green), indicating engagement of ALP. The nucleus are stained with DAPI (blue).

## Discussion

In this study, we have established a tandem fluorescent autophagy reporter system in NMR skin fibroblasts, enabling direct live-cell, flux-resolved interrogation of ALP behaviour in living NMR cells under basal conditions and during lysosomal stress. By integrating live-cell imaging, pharmacological perturbation and ultrastructural analysis, we can study time-dependent changes in the NMR autophagy pathway. Specifically, although vacuolation upon CQ-treatment has been previously observed in NMR cells using SEM approaches [20], our new tool allows us to gain spatial-temporal resolution about this process and shed light on the induction and reversibility of this phenomenon, suggesting that the phenotype reflects a regulated cellular response to lysosomal stress rather than acute cytotoxic damage. Moreover, this response is not isolated to cultured, immortalised cells and can be observed in primary NMR skin fibroblasts.

A central challenge in autophagy research is distinguishing increased autophagosome biogenesis from impaired degradation, particularly when relying on static markers such as LC3II/LC3I accumulation [9, 17, 19]. By validating the mCherry–EGFP–LC3^NMR^ reporter in NMR skin fibroblasts using both autophagy induction and lysosomal inhibition, we were able to directly visualise autophagic organisation and dynamics at single-cell resolution. Under basal conditions, NMR skin fibroblasts display an increased abundance of LC3-positive structures compared with conventional mammalian cells, accompanied by a mixed population of autophagosomes and autolysosomes. While previous studies have reported elevated basal autophagy in NMR tissues based primarily on biochemical endpoint analyses [5], our data extend these observations by showing that NMR skin fibroblasts contain a greater abundance of ALP-related vesicles at steady state, rather than enlarged individual vesicles. The coexistence of autophagosomes and autolysosomes under basal conditions suggests that NMR cells maintain a distinct steady-state organisation of the ALP, potentially reflecting a primed configuration that may facilitate rapid adaptation to stress, rather than simply constitutively elevated autophagic flux [8, 23].

Perturbation of lysosomal function using CQ reveals both expected and unexpected features of NMR autophagy. As anticipated, CQ treatment impaired autophagic degradation, resulting in the accumulation of LC3-II and mCherry⁺/EGFP⁺ puncta. Interestingly, CQ also induces the formation of many large cytoplasmic vacuoles in NMR skin fibroblasts, a phenotype not observed in HeLa cells under comparable conditions. Importantly, this vacuolation is not associated with acute cytotoxicity, as assessed by LDH release, nor did it represent irreversible cellular damage. Instead, CQ-induced vacuoles undergo progressive resolution following the removal of CQ, accompanied by reorganisation of LC3-positive structures and reacquisition of lysosomal acidity, indicating a reversible and regulated cellular response.

To confirm our interpretations with our mCherry–EGFP–LC3^NMR^ reporter cell line, we used SEM and TEM to obtain structural resolution of the nature of these vacuoles [37, 46–48]. Electron microscopy revealed that CQ-induced vacuoles are membrane-bound structures containing internal material rather than empty cavities, and that they co-exist with multiple ALP-related vesicular compartments. During recovery, these vacuoles reorganised into smaller, electron-dense, lysosome-like structures, consistent with dynamic remodelling of the lysosomal system. Complementary LysoTracker Red staining suggests that CQ-induced vacuoles are heterogeneous in their acidic properties and they progressively reacquired acidity during recovery, further supporting their functional reintegration into the ALP.

The observation that CQ-induced vacuolation occurs in both immortalised and primary NMR skin fibroblasts suggests that this phenotype reflects an intrinsic property of NMR cells. Notably, the absence of efficient recovery in HeLa cells after CQ treatment under identical conditions indicates that the recovery phenotype is not universal across mammalian cell models. While this comparison is constrained by differences in cell type and transformation status, it is consistent with the possibility that NMR skin fibroblasts possess an enhanced capacity for lysosomal remodelling after stress. While CQ is widely used as a lysosomal inhibitor and is often interpreted solely in terms of blocked autophagic flux [23, 31, 49], our findings indicate that, in NMR skin fibroblasts, lysosomal stress elicits an additional layer of cellular reorganisation that may buffer impaired degradative capacity.

Cytoplasmic vacuolisation has been described previously in several distinct biological contexts, including chemical stress, infection, senescence, and non-apoptotic cell death, but these phenomena differ substantially in mechanism and outcome [50]. Drug-induced vacuolation reported in toxicological studies is often associated with cellular injury and characterised by electron-lucent, largely empty vacuoles, with reversibility described only at a gross morphological level and without evidence of ultrastructural or functional recovery [50]. Viral infection can also induce pronounced lysosomal vacuolation, which is frequently accompanied by vacuole fusion, progressive lysosomal enlargement, and cell death [51]. In contrast, vacuolating cell death modalities such as paraptosis and methuosis are characterised by extensive ER-, mitochondrial-, or macropinosome-derived vacuolation that is LC3-independent, irreversible, and terminal [52, 53]. A distinct form of LC3-associated vacuolation has been described in oncogene-induced senescence, where enlarged vacuolar and lysosomal compartments spatially couple with LC3-positive autophagosomes to form the TOR–autophagy spatial coupling compartment (TASCC), supporting a chronic secretory phenotype [54]. Notably, this senescence-associated vacuolation is stable and irreversible, in contrast to the acute and reversible vacuolation observed in NMR skin fibroblasts. Together, these comparisons place NMR CQ-induced vacuolation in a distinct category, characterised by reversibility, lack of cytotoxicity, and dynamic reintegration into the ALP.

Our data further indicate that vacuole formation and LC3 recruitment are temporally separable processes. Vacuoles were detectable by brightfield imaging at early time points following CQ treatment, before the appearance of LC3-decorated ring structures, and the proportion of LC3-labelled vacuoles increased with prolonged exposure. This temporal dissociation suggests that LC3 recruitment is not required for initial vacuole formation. One possible interpretation is that LC3 association with CQ-induced vacuoles involves non-canonical LC3 lipidation on pre-existing single-membrane compartments (CASM) rather than classical autophagosome formation [40]. In this context, delayed LC3 recruitment and the mixed presentation of mCherry⁺/EGFP⁺ and mCherry⁺/EGFP^-^ LC3 may reflect progressive changes in vacuolar identity and luminal pH during stress adaptation, rather than canonical autophagic flux [41, 42]. While these features may be compatible with non-canonical LC3 lipidation pathways such as CASM, our data do not distinguish whether LC3 recruitment reflects CASM, canonical autophagy acting on remodelled compartments, or a mixed response triggered by lysosomal stress, and detailed investigations looking at genetic or molecular markers that distinguish CASM from canonical autophagy are still required [40, 43–45].

While we establish clear associations between CQ-induced vacuoles and ALP components, the molecular mechanisms governing vacuole formation, LC3 recruitment, and resolution remain to be defined. Moreover, CQ represents a pharmacological perturbation that may not fully recapitulate physiological lysosomal stress. Future studies employing genetic, metabolic, or *in vivo* approaches will be required to determine whether similar adaptive vacuolation responses occur under endogenous stress conditions and how they contribute to organismal resilience.

In summary, this work establishes a live-cell platform for resolving autophagy dynamics in NMR skin fibroblasts and identifies a distinctive, reversible vacuolation response to lysosomal stress. These findings support the idea that NMR cells can remodel lysosomal organisation under stress without immediately progressing to cell death, providing a framework for future mechanistic studies of autophagy–lysosome plasticity in healthy ageing. Whether similar lysosomal remodelling responses operate *in vivo* and contribute to organismal resilience or longevity remains to be determined in future physiological studies.

## Materials and Methods

### Animals

Animal experiments were performed in accordance with the United Kingdom Animal (Scientific Procedures) Act 1986 Amendment Regulations 2012 under Project Licenses (PP5814995) granted to Prof. Ewan St. John Smith by the Home Office; the University of Cambridge Animal Welfare Ethical Review Body also approved procedures. NMRs were bred in-house and maintained in an inter-connected network of cages in a humidified (∼55 %) temperature-controlled room (28–32 °C) with red lighting (08:00–16:00) and had access to food *ad libitum* and enrichment, such as chew blocks and running wheels. Animals were humanely killed by CO₂ exposure followed by decapitation.

### Reagents

All media and supplements in the following section were purchased from Gibco™ (ThermoFisher Scientific, Loughborough, UK), unless otherwise stated. Chemicals and enzymes were purchase from Merck Life Science UK Ltd. unless otherwise stated.

### Primary fibroblast isolation and culturing

Primary NMR skin fibroblast extraction was conducted on tissue isolated from NMRs sacrificed for other studies in the Smith Lab. Abdominal skin was collected from the sacrificed NMR (non-breeding male, 114w1d at death) and immediately placed in ice-cold PBS. Any residual fat or muscle tissue was carefully cleared from the skin using sterile scalpels. The skin was disinfected by spraying with an excess amount of 70 % ethanol and then washed (2X sterile PBS). The tissue was then finely minced with sterile scalpels and incubated (37°C, 3–5 h; briefly vortexing every 30 min) in an enzymatic digestion solution: 1 mg/mL collagenase, 0.2 mg/mL hyaluronidase in high-glucose DMEM. Samples were pelleted by centrifugation (2,000 rpm, 3 min), resuspended in sterile PBS (5 mL), and centrifuged again. The final pellet was resuspended in culture medium: high-glucose DMEM, 15% foetal bovine serum (Sigma), 100 U/mL penicillin-streptomycin (Gibco), 1 X MEM non-essential amino acids, 1 mM sodium pyruvate, 100 μg/mL Primocin (InvivoGen) and 100 μg/mL Normocin (InvivoGen). The cell suspension was passed through a Falcon 70-μm cell strainer (Fisher Scientific) and seeded in a treated T-75 culture flask (Greiner Bio-One, 658175). NMR cells were cultured at 32°C in a 5% CO_2_, 3% O_2_ incubator until 80% confluency was reached. All fibroblasts were used within the first 5 passages. Cells were plated in six-well dishes (2 x 10^5^ cells/well) and incubated overnight before treatment with CQ.

### Immortalised NMR skin fibroblasts

NMR skin fibroblasts were immortalised by SV40 transfection and previously characterised [32]. All cell lines were cultured as a monolayer in complete medium: high-glucose DMEM, 15% foetal bovine serum, 100 U/mL penicillin-streptomycin (Gibco), 1 X MEM non-essential amino acids, 1 mM sodium pyruvate and 100 μg/mL Primocin, in a 75 cm² tissue culture flask with a filter cap (Greiner Bio-One) and incubated in a humidified 32°C incubator, supplied with 5% CO₂ and 3% O₂ atmosphere. Cell passaging was performed when the cells reached 80–90% confluence. The culture medium was removed, and the cell monolayer was washed (1X sterile PBS (5 mL)). Cells were detached using 1× trypsin–EDTA, prepared from a 10× stock solution, followed by the addition of complete medium to neutralise trypsin. The cell suspension was centrifuged (500 ×g, 5 min, room temperature (RT)), and the cell pellet was resuspended in 1 mL fresh culture medium. Cell viability and density were determined using a haemocytometer and trypan blue exclusion staining (Gibco), and the cells were seeded (2 × 10⁶ cells in 10 mL fresh media, T-75 flask). The medium was changed every 3–4 days, and all experiments were conducted using cells between passages 3-20.

### Establishing the mCherry–EGFP–LC3^NMR^ reporter NMR skin fibroblast cell line Plasmids

The pDEST-CMV mCherry–EGFP–LC3^WT^ human plasmid expressing an mCherry–EGFP–LC3 fusion protein was a gift from Robin Kettler (Addgene plasmid #123230) [55]. The amino acid sequence of human and NMR LC3 was BLAST aligned (https://blast.ncbi.nlm.nih.gov/Blast.cgi (Supplementary Table 1) and primers were designed (Supplementary Table 2) to modify the human LC3 into an NMR version by introducing E105G, M121R and K122G. Mutations were introduced using the QuikChange Site-Directed Mutagenesis protocol (Agilent Technologies LDA UK Ltd.), and the plasmid was purified using a Qiagen Plasmid Midi kit (Qiagen). pDEST-CMV mCherry–EGFP–LC3^NMR^ was confirmed by DNA sequencing.

### pDEST-CMV mCherry–EGFP–LC3^WT^ expressing HeLa cell line

HeLa Kyoto cells were cultured in Dulbecco’s Modified Eagle’s Medium (DMEM) + GlutaMAXTM (Gibco) with 10% Fetal Bovine Serum. A killing curve for HeLa Kyoto cells with a range of G418 Sulfate (Gibco) concentrations was made to determine 600 µg/ml as the selective concentration. HeLa Kyoto cells were transfected with 20 µg mCherry–EGFP–LC3^WT^ DNA (Addgene Plasmid #123230), using the Neon NxT Electroporation System (ThermoFisher) according to the manufacturer’s instructions. After 48 hr the cells were transferred onto 10 cm dishes at a range of dilutions, with 600 µg/ml G418 Sulfate. The media on the dishes was changed every 3–4 days. Distinct colonies were selected and transferred to 12-well plates to grow. Clones were screened by Western blot to confirm target protein expression and further validated with microscopy to confirm the desired functional response.

### pDEST-CMV mCherry–EGFP–LC3^NMR^ expressing NMR skin fibroblast cell line

G418 sulphate (100 mg/mL stock) was used as a selective agent to establish cell lines expressing pDEST-CMV mCherry–EGFP–LC3^NMR^ based on neomycin resistance. The resistance of non-transfected immortalised NMR skin fibroblast cells was determined by treatment with different G418 sulphate concentrations. It was established that 4 mg/mL G418 sulphate resulted in 80% cell death and this concentration was used for further selection. Immortalised NMR skin fibroblasts were transfected using FuGENE® HD Transfection Reagent (Promega UK) with a 1:3 DNA-to-reagent (w/v) ratio (2.5 µg plasmid DNA, 7.5 µL reagent), following the manufacturer’s protocol and plated in a 6-well cell culture plate until reaching 60-70% confluency. Cells were trypsinised and plated in 10-cm cell culture dishes and after 48 h, G418 sulphate was added, and the transfected cells were cycled between selection media (7-day) and non-selection media (4-day) until desired single colony growth was established. Established colonies were cultured in 12-well cell culture plates in the presence of G418 sulphate and the resulting cell lines were characterised.

### Western Blotting (WB)

Cells were plated in a 6-well plate and trypsinised when confluency reached 80-90%. The cells were resuspended in 1 mL PBS and centrifuged (500 × g, 5 min, RT), followed by resuspension (with gentle pipetting) in 125 μL ice-cold lysis buffer (10 mL bioPLUS RIPA buffer (pH 8) (BioWorld, USA), one tablet of cOmplete™ Mini Protease Inhibitor Cocktail (Roche Diagnostics), and 1.5 μL BaseMuncher Endonuclease (Abcam)) and incubated (20 min, on ice). The cells were mixed with gentle pipetting (6-8X) and further incubated (10 min, on ice). Cell lysates were centrifuged (12,000 rpm, 4°C, 12 min), and the supernatants and Spectra^TM^ Multicolour Broad Range Protein Ladder (ThermoFisher Scientific) were separated on 15% SDS-PAGE gels (1.5 mm thickness) in 1X Tris-glycine-SDS (TGS) running buffer (60V (30 min), 120V (90 min)). Proteins were transferred onto Immobilon-P polyvinylidene difluoride (PVDF) membranes (Merck Millipore) using a Pierce™ Power Blot Cassette system (25 V, 1.3 A, 30 min). The membranes were blocked in 5% (w/v) dried skimmed milk prepared in Tris-buffered saline with 0.1% Tween (TBS-T) (1 h, RT, gentle agitation) followed by 3X washes with TBS-T (5 min). The membranes were cut according to target molecular weight (using the protein ladder as a guide) and incubated separately (4°C, overnight (ON)) with an appropriate primary antibody prepared in 5% bovine serum albumin (BSA) in TBS-T supplemented with 0.02% sodium azide (rabbit anti-LC3B primary antibody (GeneTex, GTX127375, 1:1000 dilution), rabbit anti-GAPDH primary antibody (Proteintech, 10494-1-AP, 1:1000 dilution), or mouse anti-GFP primary antibody (Takara Bio, 632380, 1:1000 dilution)).

Membranes were washed in TBS-T (3X, 5 min) and incubated (1 h, RT) with a horseradish peroxidase-conjugated secondary antibody (swine anti-rabbit IgG (Dako, P0399, 1:5000 dilution) for anti-LC3B and anti-GAPDH; rabbit anti-mouse IgG (Dako, P0260, 1:5000 dilution) for anti-GFP prepared in 5% skimmed milk/TBS-T). After TBS-T washing (3X, 5 min), blots were incubated with Amersham™ ECL Western Blotting Detection Reagent (Cytiva) and imaged using an Odyssey® Fc Imaging System (LI-COR Biosciences) with exposure times of 30 s to 2 min. Image acquisition was performed using 90% brightness and 80% contrast settings.

### Immunofluorescence

Cells were plated on sterile glass coverslips (VWR International) in 6-well plates and cultured until 70% confluency was reached. Culture medium was removed and cells were washed with PBS (3X, 5 min), followed by the addition of 4% formaldehyde in PHEM buffer (60 mM PIPES, 25 mM HEPES, 10 mM EGTA, 4 mM MgSO_4_; 2 mL per well) and the cells were incubated (12 min, RT). The cells were washed with PBS (3X, 5 min) and incubated in 5% BSA in TBS-T (1 h, RT). After washing with PBS (3X, 5 min), the cells were incubated with rabbit anti-LC3B primary antibody (GeneTex, 1:600 dilution) in PBS-T (4°C, ON), washed with PBS (3X, 5 min) and incubated with goat anti-rabbit IgG (H+L) cross-adsorbed Alexa Fluor™ 488-conjugated secondary antibody (ThermoFisher Scientific, A-11008, 1:300 dilution). After final washes in PBS (3X, 5 min), coverslips were mounted using VECTASHIELD® Antifade Mounting Medium with DAPI (2BScientific, H-1200) and imaged on a STELLARIS 5 confocal microscope (Leica Microsystems) following manufacturer-recommended settings, at the Microscopy Bioscience Platform, University of Cambridge, UK.

### Pharmacological Treatments

PP242 (Selleck Chemicals, S2218) and bafilomycin A1 (Cayman Chemical, 11038) were prepared as DMSO stock solutions. Chloroquine diphosphate salt (Sigma-Aldrich, C6628) was dissolved in sterile water. Cells were treated with PP242 (2–4 µM, 24 h), bafilomycin A1 (166–333 nM, 24 h), or chloroquine (40–100 µM, 4–24 h). Control cells received equivalent volumes of DMSO. For recovery experiments, chloroquine-containing media was removed, cells were washed (1X PBS) and fresh media was added for recovery periods of 4 h or 24 h.

### LysoTracker Red Staining

Cells were incubated with LysoTracker™ Red DND-99 (ThermoFisher Scientific, L7528) at a final concentration of 75 nM (32°C, 12 min) prior to live-cell imaging. Following incubation, cells were washed (1X) with pre-warmed media and imaged immediately.

### Live-Cell Imaging and Image Analysis

Live-cell imaging of mCherry–EGFP–LC3^NMR^ reporter cells was performed at 32°C and 5% CO_2_ under atmospheric O₂ conditions, and mCherry–EGFP–LC3^WT^ HeLa reporter cells was performed at 37°C and 5% CO_2_ under atmospheric O₂ conditions, both using a Leica STELLARIS 5 confocal microscope. EGFP and mCherry fluorescence signals were acquired sequentially using identical acquisition settings across experimental conditions. Brightfield images were acquired in parallel to visualise cellular morphology and vacuole formation. Quantitative analysis of LC3-positive structures was performed using Fiji [56]. Cell area was manually defined, LC3 puncta were identified using consistent thresholding parameters, and puncta number was normalised to cell area. Puncta diameter was measured using the diameter tool. Statistical analyses were performed as specified in the figure legends.

### Electron microscopy

Electron microscopy was performed with support from the EM facility of the Microscopy Bioscience Platform of the School of Biological Sciences, University of Cambridge, UK. Immortalised NMR skin fibroblasts were either untreated, treated with CQ (100 µM) for 24 h, or allowed to recover in fresh media for 24 h following CQ removal. Samples were fixed in 2% glutaraldehyde and 2% formaldehyde in 0.05 M sodium cacodylate buffer (pH 7.4) supplemented with 2 mM calcium chloride (4°C, ON). After fixation, samples were washed (5X, 0.05 M sodium cacodylate buffer (pH 7.4)) subsequently osmicated using a solution containing 1% osmium tetroxide and 1.5% potassium ferricyanide in 0.05 M sodium cacodylate buffer (pH 7.4) (4°C, 3 days). Samples were then washed in deionised water (DIW) (5X) and incubated with 0.1% (w/v) thiocarbohydrazide in DIW (20 min, RT, avoid of light). Following 5X DIW washes, samples were osmicated a second time using 2% osmium tetroxide in DIW (1 h, RT). After further washing in DIW (5X), samples were block-stained with 2% uranyl acetate in 0.05 M maleate buffer (pH 5.5) (4°C, 3 days). Following staining, samples were washed with DIW (5X) and dehydrated through a graded ethanol series (50%, 70%, 95%, 100%, and 100% dry ethanol), followed by 100% dry acetonitrile (3X 5 min wash each solution). Samples were then infiltrated with a 1:1 mixture of 100% dry acetonitrile and Quetol resin (without BDMA) overnight, followed by incubation in 100% Quetol resin (without BDMA) for 3 days. Subsequently, samples were infiltrated in 100% Quetol resin containing BDMA for 5 days, with the resin exchanged daily. The Quetol resin mixture consisted of 12 g Quetol 651, 15.7 g NSA (nonenyl succinic anhydride), 5.7 g MNA (methyl nadic anhydride), and 0.5 g BDMA (benzyldimethylamine) (all from TAAB). Samples were polymerised at 60°C for 2 days.

For SEM block-face imaging, semi-thin sections (∼200 nm) were cut using an ultramicrotome (Leica Ultracut) and mounted on Melinex plastic coverslips, where they were allowed to air-dry. Coverslips were mounted on aluminium SEM stubs using conductive carbon tabs, and the edges were painted with conductive silver paint. Samples were then sputter-coated with a 30 nm carbon layer using a Quorum Q150 T E carbon coater. Imaging was performed using a Verios 460 scanning electron microscope (FEI/Thermo Fisher Scientific) operated at an accelerating voltage of 4 keV and a probe current of 0.2 nA in backscatter mode using the concentric backscatter detector (CBS). Field-free mode was used for low-magnification imaging, while immersion mode was used for high-magnification imaging. Images were acquired at a working distance of 3.5–4 mm with a resolution of 1536 × 1024 pixels, a dwell time of 3 µs, and four-line integrations. Stitched maps were generated using FEI MAPS software with the default stitching profile and 5% image overlap.

For TEM imaging, ultrathin sections (∼70 nm) were cut and mounted on bare copper TEM grids. Sections were imaged using a Tecnai G2 transmission electron microscope (FEI/Thermo Fisher Scientific) operated at 200 keV with a 20 µm objective aperture to improve contrast. Images were acquired using a high-resolution CCD camera (Advanced Microscopy Techniques Corp., Danvers, USA).

### Cytotoxicity assay

Cytotoxicity was assessed using the CytoTox 96® Non-Radioactive Cytotoxicity Assay (Promega). Cells were seeded into 96-well Cell Culture Plates (Corning) at a density of 1 × 10⁴ cells per well and allowed to adhere for 24 h prior to treatment. Following experimental treatments, and immediately before the assay, 10 μL of 10× Lysis Solution was added to designated wells to generate maximum LDH release controls and incubated (37°C, 45 min).

Culture media were collected, and 50 μL from each well was transferred to a fresh 96-well assay plate, followed by addition of 50 μL CytoTox 96® Reagent. Plates were incubated protected from light (30 min, RT). The reaction was terminated by addition of 50 μL Stop Solution, and absorbance was measured at 490 nm using a FLUOstar® Omega microplate reader (BMG LABTECH).

Percentage cytotoxicity was calculated as the following:

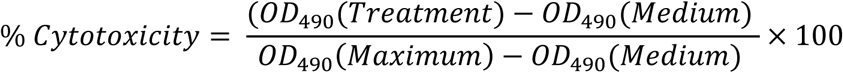

### Statistical analysis

All experiments were performed with at least three independent biological replicates unless otherwise stated. Data are presented as mean ± SEM. Statistical analyses were conducted using GraphPad Prism 9 (GraphPad Software, San Diego, CA, USA), with specific tests indicated in the corresponding figure legends.

## Acknowledgements

The authors would like to thank Dr. Jonathan Howe (Microscopy Bioscience Platform, University of Cambridge) for advice on confocal microscopy imaging. All electron microscopy imaging was performed at the EM facility of the Microscopy Bioscience Platform, University of Cambridge. F.T., M.P.H, E.S.S. and J.R.K. acknowledge support from the David James Trust Fund and M.P.H is partially supported by a Cambridge Philosophical Society Studentship. This work was supported by an MRC Career Development Award (MR/W01632X/1, J.R.K.), the Royal Society (RGS\R1\231207, J.R.K.) and the Leona M. & Harry B. Helmsley Charitable Trust (2508-08462, E.S.S.).

