## Supplementary Information for "A live-cell autophagy reporter reveals reversible vacuolation in naked mole-rat skin fibroblasts under lysosomal stress"

Fangze Tong[1], Mateo P. Hoare[1], Laura J. Grundy[1], Filomena Gallo[2], Karin H. Müller[2], Ewan St. John Smith[1*] and Janet R. Kumita[1*]

[1]Department of Pharmacology, University of Cambridge, Tennis Court Road, Cambridge, CB2 1PD, UK.

[2]Elecron Microscopy Facility of the Microscopy Bioscience Platform of the School of Biological Sciences, University of Cambridge, Anatomy Building, Downing Site, Cambridge CB2 3DY, UK.

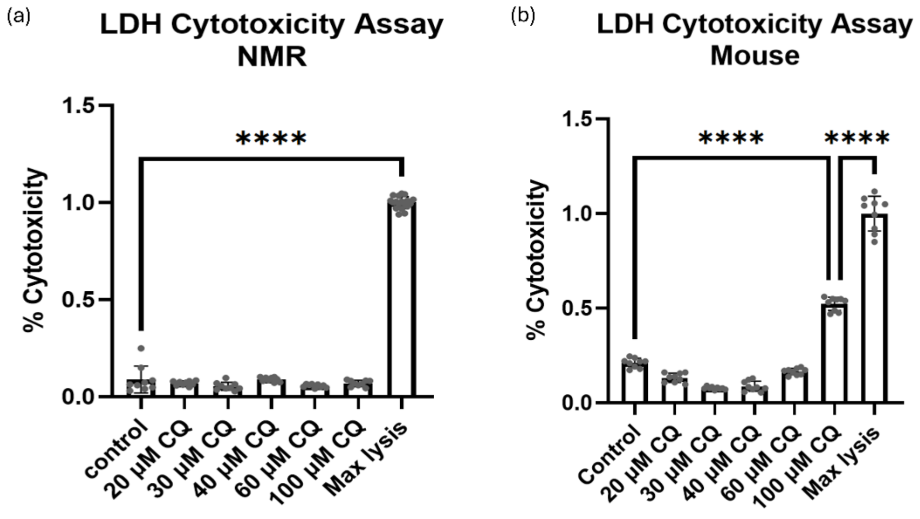


**Supplementary Figure 1: Percent cytotoxicity of immortalised NMR (A) and mouse (B) skin fibroblast**. An LDH-based cytotoxicity assay was performed on NMR and mouse skin fibroblasts (immortalised using the same method as the NMR skin fibroblasts) [1]. Both NMR and mouse skin fibroblasts were either untreated or treated with CQ for 24 h (20, 30, 40, 60, and 100 µM). Results from each experiment were standardised to maximum lysis with Triton X, pooled (n=9), and analysed using one-way ANOVA. ****, $p<0.001$. The results showed that even at the highest CQ concentration (100 µM), treated NMR skin fibroblasts exhibited no significant cytotoxicity.

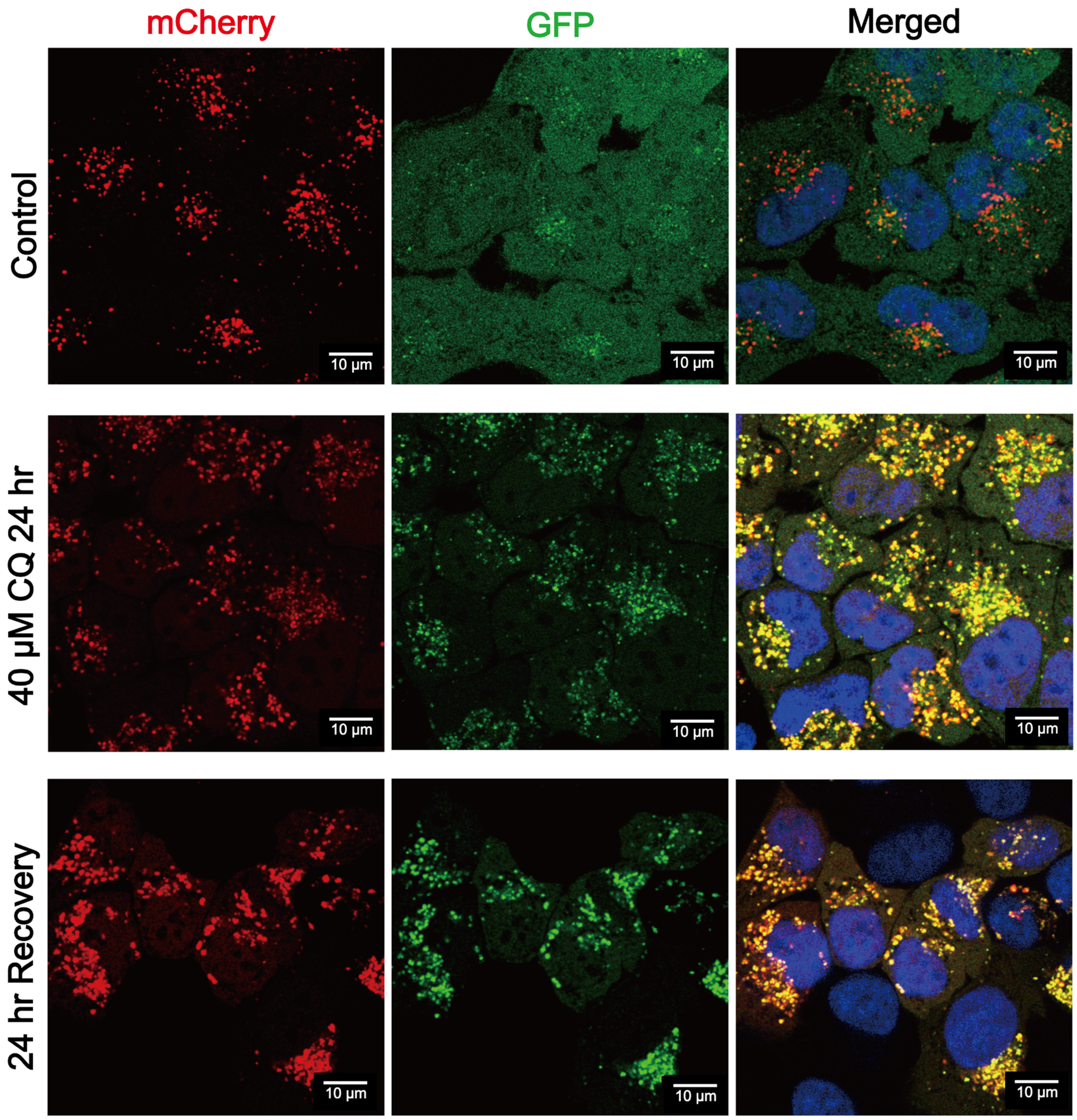


**Supplementary Figure 2. Limited recovery of autophagic organisation following CQ removal in HeLa cells.** Representative live-cell confocal images of HeLa cells stably expressing the tandem fluorescent autophagy reporter mCherry–EGFP–LC3$^{WT}$. Cells were either left untreated, treated with CQ (40 µM) for 24 h, or allowed to recover for 24 h following CQ removal. The nucleus is stained with Hoescht (Blue). After 24 h of CQ treatment, LC3-positive structures predominantly appear as mCherry$^{+}$/EGFP$^{+}$ (yellow) puncta. In contrast to NMR skin fibroblasts, removal of CQ does not result in substantial reorganisation of LC3-positive structures, as the majority of LC3 puncta remain mCherry$^{+}$/EGFP$^{+}$ (yellow) after 24 h of recovery. All images were acquired using identical imaging settings.

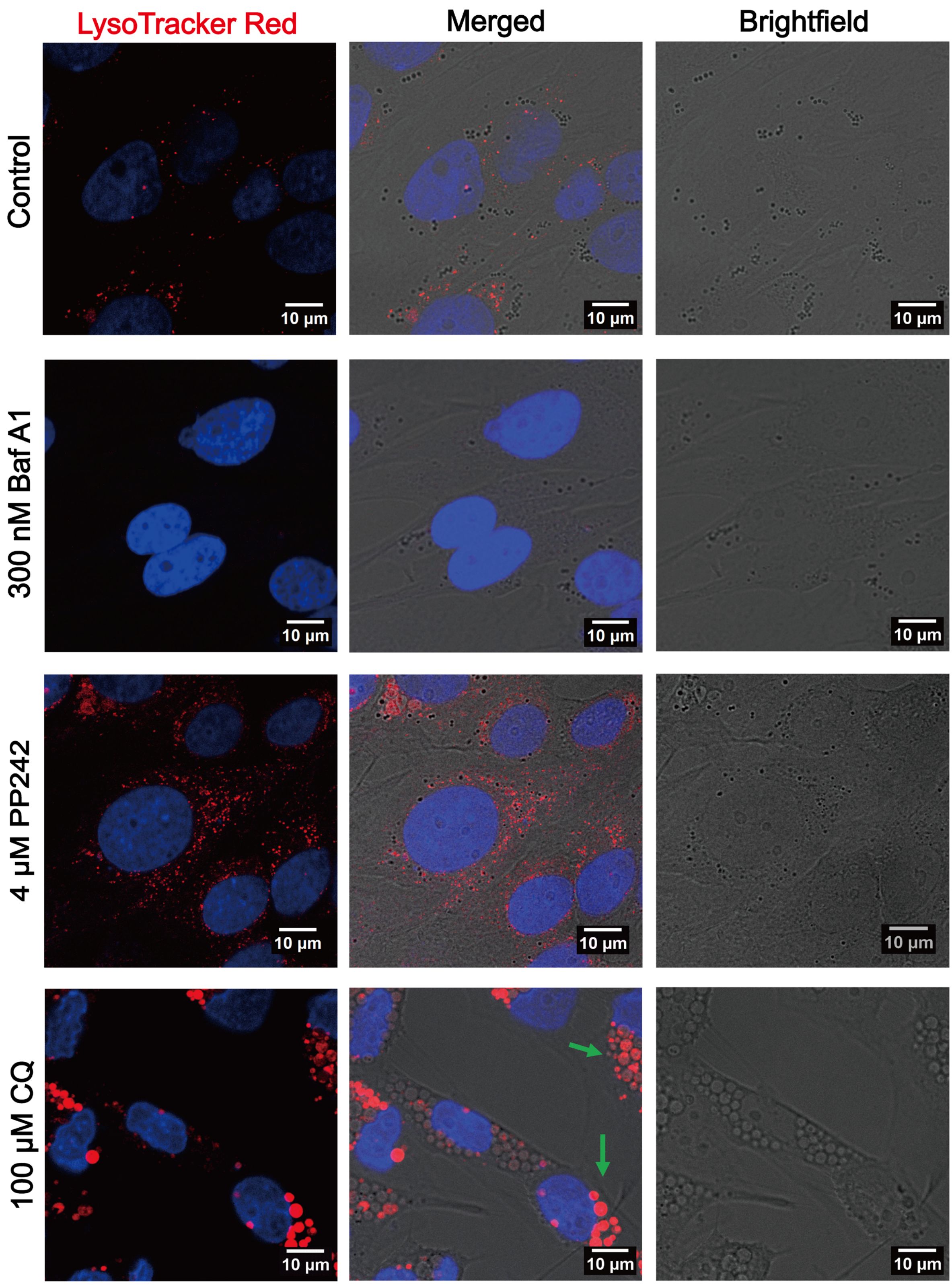


**Supplementary Figure 3: Characterisation of LysoTracker Red staining in immortalised NMR skin fibroblasts.** Representative live-cell confocal images of immortalised NMR skin fibroblasts stained with LysoTracker Red and Hoescht (nucleus – blue). Cells were either untreated or treated (200 nM Baf A1, 4 µM PP242, or 100 µM CQ) for 24 h. In Baf A1-treated cells, LysoTracker Red signal was completely abolished as Baf A1 is a vacuolar proton pump inhibitor that disrupts lysosomal pH. In contrast, PP242 treatment enhanced LysoTracker Red signalling due to upregulation of autophagy upon mTOR inhibition. Interestingly, in CQ-treated NMR skin fibroblasts, the overall LysoTracker Red signal was more prominent, but the staining was not uniformly associated with all vacuoles, and only some vacuoles were strongly stained by LysoTracker Red (green arrows).

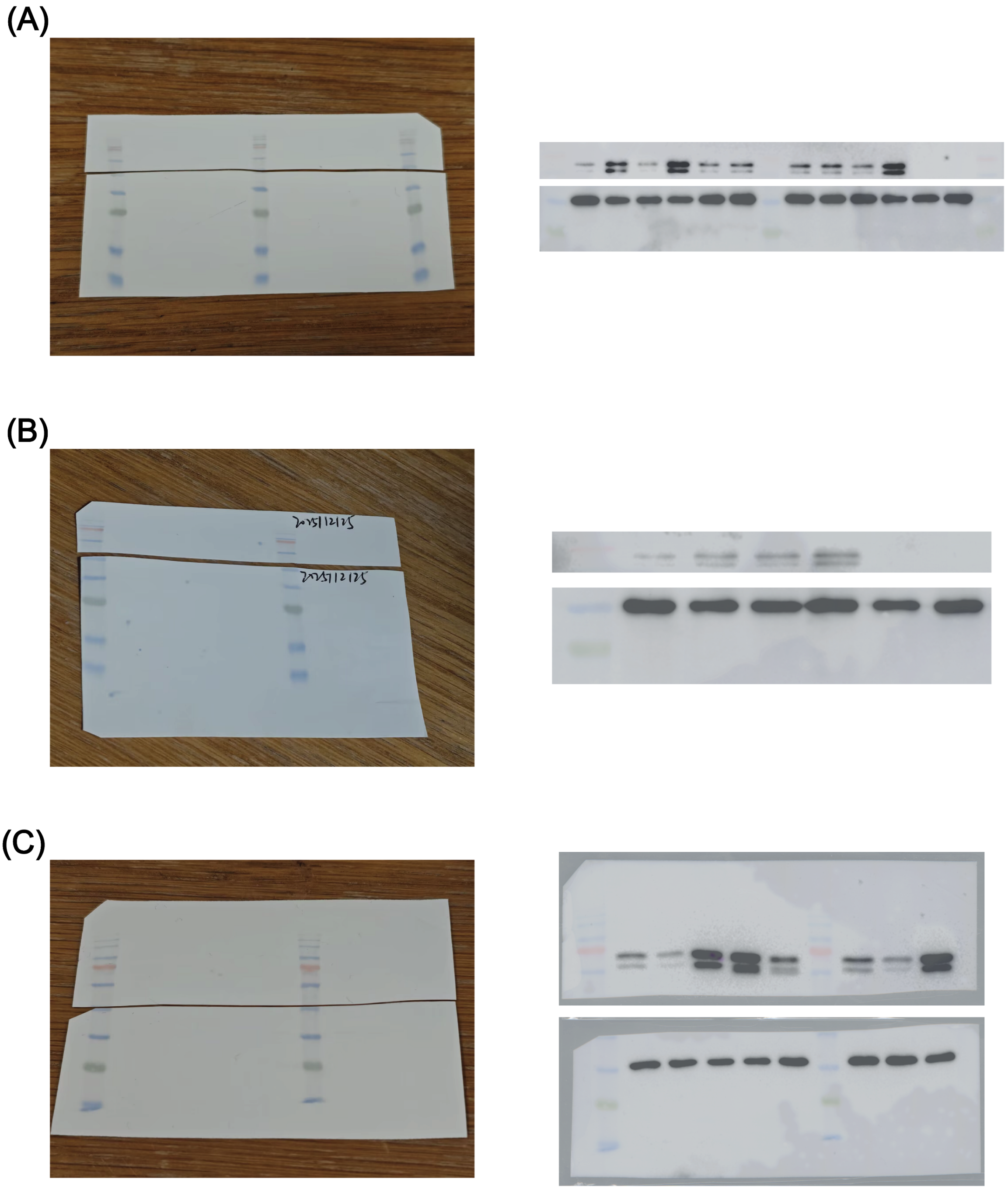


**Supplementary Figure 4: Full western blot membranes. A)** Full membrane and blot correspond to Figure 1A. Originally, 12 colonies were characterised, and colony 2 was taken forward. **B)** Full membrane and blot correspond to Figure 1B. **C)** Full membrane and blot correspond to Figure 3B. The three lanes on the very right side were technical replicates of the same samples.

**Supplementary Table 1: Protein sequence alignment of the NMR and human LC3.** NMR and human LC3 were aligned using BLASTn, and differences in amino acid sequence are highlighted in yellow.

| **Score** | **Except** | **Method** | **Identities** | **Positives** | **Gaps** |
|---|---|---|---|---|---|
| 248 bits (634) | 4e-92 | Compositional matrix adjust. | 122/125 (98%) | 122/125 (97%) | 0/125 (0%) |

| | | |
|---|---|---|
| **NMR LC3** | MPSEKTFKQRRTFEQRVEDVRLIREQHPTKIPVIIERYKG | 40 |
| | MPSEKTFKQRRTFEQRVEDVRLIREQHPTKIPVIIERYKG | |
| **Human LC3** | MPSEKTFKQRRTFEQRVEDVRLIREQHPTKIPVIIERYKG | 40 |
| **NMR LC3** | EKQLPVLDKTKFLVPDHVNMSELIKIIRRRLQLNANQAFF | 80 |
| | EKQLPVLDKTKFLVPDHVNMSELIKIIRRRLQLNANQAFF | |
| **Human LC3** | EKQLPVLDKTKFLVPDHVNMSELIKIIRRRLQLNANQAFF | 80 |
| **NMR LC3** | LLVNGHSMVSVSTPISEVYESEKDGDGFLYMVYASQETFG | 120 |
| | LLVNGHSMVSVSTPISEVYESEKD DGFLYMVYASQETFG | |
| **Human LC3** | LLVNGHSMVSVSTPISEVYESEKDEDGFLYMVYASQETFG | 120 |
| **NMR LC3** | RGLSV | 125 |
| | LSV | |
| **Human LC3** | MKLSV | 125 |

**Supplementary Table 2: Primers used for SDM to create pDEST CMV mCherry-EGFP-LC3$^{NMR}$.**

| Name | Purpose | Sequence |
|---|---|---|
| E105G Forward | SDM of the pDEST-CMV mCherry–EGFP–LC3$^{NMR}$ plasmid | GTATGAGAGTGAGAAAGATGGAGATG GATTCCTGTACATGG |
| E105G Reverse | SDM of the pDEST-CMV mCherry–EGFP–LC3$^{NMR}$ plasmid | CCATGTACAGGAATCCATCTCCATCTT TCTCACTCTCATAC |
| M121R Forward | SDM of the pDEST-CMV mCherry–EGFP–LC3$^{NMR}$ plasmid | GTTCGGGCGGAAATTGTCAGTGTAAT ACCCAGCT |
| M121R Reverse | SDM of the pDEST-CMV mCherry–EGFP–LC3$^{NMR}$ plasmid | AATTTCCGCCCGAACGTCTCCTGGGA |

| | | |
|---|---|---|
| K122G Forward | SDM of the pDEST-CMV mCherry–EGFP–LC3$^{NMR}$ plasmid | GGGTATTACACTGACAATCCCCGCCC GAACGTCTCCT |
| K122G Reverse | SDM of the pDEST-CMV mCherry–EGFP–LC3$^{NMR}$ plasmid | AGGAGACGTTCGGGCGGGGATTGTCA GTGTAATACCC |
